# A Multiresolution Approach with Method-Informed Statistical Analysis for Quantifying Lymphatic Pumping Dynamics

**DOI:** 10.1101/2024.04.24.590950

**Authors:** Mohammad S. Razavi, Katarina J. Ruscic, Elizabeth G. Korn, Marla Marquez, Timothy T. Houle, Dhruv Singhal, Lance L. Munn, Timothy P. Padera

**Affiliations:** Edwin L. Steele Laboratories, Department of Radiation Oncology, Massachusetts General Hospital Cancer Center, Massachusetts General Hospital and Harvard Medical School, Boston, MA 02114, USA; Massachusetts General Hospital Department of Anesthesia, Critical Care and Pain Medicine and Harvard Medical School, Boston, MA 02114, USA; Beth Israel Deaconess Medical Center, Division of Plastic and Reconstructive Surgery, Department of Surgery, Boston, MA, 02215, USA

## Abstract

Despite significant strides in lymphatic system imaging, the timely diagnosis of lymphatic disorders remains elusive. One main cause for this is the absence of standardized, quantitative methods for real-time analysis of lymphatic contractility. Here, we address this unmet need by combining near-infrared lymphangiography imaging with an innovative analytical workflow. We combined data acquisition, signal processing, and statistical analysis to integrate traditional peak and-valley with advanced wavelet time-frequency analyses. Decision theory was used to evaluate the primary drivers of attributable variance in lymphangiography measurements to generate a strategy for optimizing the number of repeat measurements needed per subject to increase measurement reliability. This approach not only offers detailed insights into lymphatic pumping behaviors across species, sex and age, but also significantly boosts the reliability of these measurements by incorporating multiple regions of interest and evaluating the lymphatic system under various gravitational loads. By addressing the critical need for improved imaging and quantification methods, our study offers a new standard approach for the imaging and analysis of lymphatic function that can improve our understanding, diagnosis, and treatment of lymphatic diseases. The results highlight the importance of comprehensive data acquisition strategies to fully capture the dynamic behavior of the lymphatic system.

## INTRODUCTION

The lymphatic system is vital for the maintenance of tissue fluid balance and immune integrity in the human body[1]. Despite this, our understanding of lymphatic function has been neglected relative to other organ systems. Lymphedema is a lymphatic disease marked by abnormal accumulation of lymph fluid in tissues. Lymphedema affects one in 1000 Americans and leads to infections, chronic pain, loss of limb function, and lymphangiosarcoma[2, 3]. Monitoring of lymphatic function and treatment response in patients with lymphatic diseases is hindered by a reliance on using disease signs for diagnosis. At present, lymphatic diseases such as lymphedema cannot be diagnosed until the disease has progressed enough to develop early clinical signs, such as increased limb volume. While several methods such as lymphoscintigraphy, magnetic resonance imaging (MRI) lymphangiography, x-ray lymphography, fluorescein microlymphangiography and near infrared fluorescence (NIRF) lymphangiography are available at specialized medical centers to evaluate lymphatic anatomy and rudimentary lymphatic function, there are no robust, standardized techniques for imaging real-time lymphatic contractility used in current standard medical practice[4]. While NIRF lymphangiography is the newest of these methods and has the most promise for measuring lymphatic contractility, there is both a lack of standardization among different camera systems[5], and a lack of standardized analysis methods to optimize the functional data derived from the video data obtained with this technique. The lack of diagnostic capability for lymphatic diseases and visualization of the lymphatic system prevents the timely diagnosis of lymphatic disease and impedes research for the development of novel treatments and routine monitoring of lymphatic function. Thus, there is a pressing need for improved lymphatic imaging and quantification methods to enable prompt lymphedema diagnosis and treatment, which has prompted ARPA-H to make overcoming this clinical deficiency a priority.

Collecting lymphatic vessels are comprised of tightly interconnected lymphatic endothelial cells (LECs) and specialized lymphatic muscle cells (LMCs). The contractile unit of lymphatic vessels—the lymphangion—is the region between intraluminal valves that pump as a result of spontaneous contractions of LMCs[6, 7]. The unidirectional valves counteract lymph backflow when there is an opposing pressure gradient, which is affected by body positioning relative to gravity[8, 9]. Unlike the cardiovascular system, in which intravenously introduced contrast travels to the heart, and is circulated through the entire cardiovascular tree, the unidirectional valves and distal-to-central flow of the lymphatic system mean that a central injection does not backfill and illuminate the lymphatic network in normal conditions, though dermal backflow can be observed in abnormal states[10]. Lymphatic imaging modalities have evolved to take advantage of interstitial or intranodal injection of tracers, which then travel to local collecting lymphatic vessels[11]. Measurement and quantification of lymphatic contraction remains challenging, prompting ongoing evolution of innovative methods to characterize this fundamental, physiological vascular system.

Traditional lymphoscintigraphy, while considered the gold standard for lymphedema diagnosis, involves patient exposure to ionizing radiation, requires specialized imaging facilities and personnel, and affords only limited spatial resolution of the local lymphatic network[12–14]. This method offers some insight into lymphatic function, as decreased uptake or characteristic distributions of tracer at the endpoint of the study can occur in advanced disease. However, early lymphedema is not readily detected[15]. The approach does not rely on real-time measurement of lymphatic pumping, but rather pre and post-snapshots of lymphatic network filling to calculate a lymph transit time, from which a large decrease in function can be inferred[4]. While x-ray lymphography and magnetic resonance lymphography afford much higher spatial anatomic resolution, like lymphoscintigraphy, they do not provide information about real-time lymphatic contractile function. Fluorescence imaging with visible light wavelengths *in vivo* is limited by signal-to-noise attenuation from tissue scattering, absorption, and autofluorescence, which prevents imaging of structures more than a few millimeters deep. In preclinical models, this depth limitation can be somewhat bypassed by removing the skin overlying the area being studied[16]. Removal of the skin is not translatable to a human bedside diagnostic tool.

NIRF lymphography, which utilizes excitation light greater than 750 nm, allows for greater depth of tissue imaging (3-4 cm) with less autofluorescence compared to fluorescence imaging with visible wavelengths and does not require skin removal[17, 18]. Studies from 2005-2008 first reported its use in humans and describe the injection of indocyanine green (ICG) intradermally, which spreads locally to collecting lymphatic vessels in an anatomic region, such as a single limb[19–22]. While the use of ICG-based NIRF lymphography has expanded, it remains used primarily for anatomic mapping in humans[23]. NIRF technology and analysis has also been developed and investigated in preclinical models[24], primarily in mice and rats. Lymphatic contractility has been quantified via NIR lymphography by plotting signal intensity in a small region of interest (ROI) over time. These studies have been used for many wide-ranging, exciting applications, including detecting differences in lymphatic contraction rates between wild-type mice and those bearing B16 lymph node metastases[25], untreated lymphatics in rats and those exposed to topical nitric oxide[26], normal and high salt diet in mice and rats[27], and swine getting manual lymphatic drainage compared to baseline[28].

A major challenge in the application of these techniques is choosing the region of interest (ROI) for analyzing the contractions. The assignment of ROIs remains a subjective manual process and there is no standardized method concerning the placement of ROIs for analyses, making it potentially difficult to compare results from different studies. Pulsatile lymph flow and the movement of “packets” of NIR dye in the lymphatic vessels lead to increasing and decreasing fluorescent intensity over the time-scale of a contraction. The resulting time series data can be analyzed by identifying local minima and maxima (termed peaks and valleys). The evaluation of the frequency and amplitude of signal peaks is the current state-of-the-art in the lymphatic field for extrapolating information on lymphatic contraction frequency and strength in preclinical models[25, 26, 29].

Lymphatic research has taken steps towards quantifying human NIRF lymphangiography, and several groups have applied a similar approach to human NIRF lymphangiography and quantification, but with mixed repeatability[30–32]. For this reason, there is a pressing need for ongoing improvement in analysis methods for the quantification of lymphatic contractility measured by NIRF lymphangiography in preclinical models. This would enable translation to human studies using similar methods.

In this study, we designed a workflow and developed quantitative tools to capture the different components that describe lymphatic function measured by NIRF lymphangiography. In our workflow, we consider multiple ROIs, exposing mice to various positions and loading conditions to unveil differences in lymphatic pumping behavior in both adult and aged mice of different sexes. We analyze signals using traditional peak and valley analysis, as well as multi-resolution wavelet-based techniques, to evaluate lymphatic pumping. The use of several ROIs is crucial for being able to make multiple measurements simultaneously from a single subject and increases the reliability of the measured lymphatic contractility. We also designed linear mixed models to account for systematic and intergroup variabilities. Our technique suggests that exposing lymphatic vessels to different loading conditions, along with using multiple ROIs, better reveals nuances in lymphatic pumping function. Our image processing and statistical methods were also applied to human NIRF data.

## RESULTS

### Workflow for data acquisition, signal processing, and statistical analysis for improved quantification of lymphatic pumping

One limitation of current methods, including the peak and valley method, is that irregular contraction patterns are not distinguished from regular contraction rhythms of the same time-averaged frequency. This limitation can introduce variability in ultimate frequencies measured when only short recordings are analyzed and also ignores what are likely important physiological states of lymphatic contractility. Our study proposes that an essential concept to make *in vivo* lymphangiography a technique that can give meaningful, reliable results, with the ultimate goal of human translation, is to collect repeated measurements from the same subject, to measure instantaneous frequencies and other contractility parameters, and to study the system in different states of behavior. Dynamic physiological systems need to be studied in a range of states to capture their essential properties. We chose gravitational stress as the impulse response system to explore the range of behavior of the lymphatic vessel network. In previously collected human ICG lymphangiography and murine intravital epifluorescence lymphangiography, data from different gravitational stresses was not available, but the power of repeated measurements and time-frequency analysis were also shown using our methods.

For mouse NIRF lymphangiography gravitational studies, a custom-built frame around the NIRF imaging apparatus allowed for complete positioning of the mouse in different orientations while keeping the distance between the lymphatic vessels and excitation source and emission collection instrumentation exactly the same (Figure 1A). For all data streams, movies were processed in MATLAB using custom code. ROIs were selected and denoised. In each ROI, the signal intensity was plotted with respect to time, creating a “peak and valley plot.” Local minima and maxima were identified, and frequency was calculated by the simple division of cycles (or peaks) per unit time, while amplitude was the total height of the signal from a maximum to a minimum. Note that this also relies on a user-selected threshold for amplitude minima for the peaks to distinguish a true contraction from the noise in the data in the calculation of frequency. This is the traditional method by which intravital NIRF has been analyzed, often with a single or few subjectively "best" ROI(s) chosen by the experimenter for analysis.

**Figure 1:**
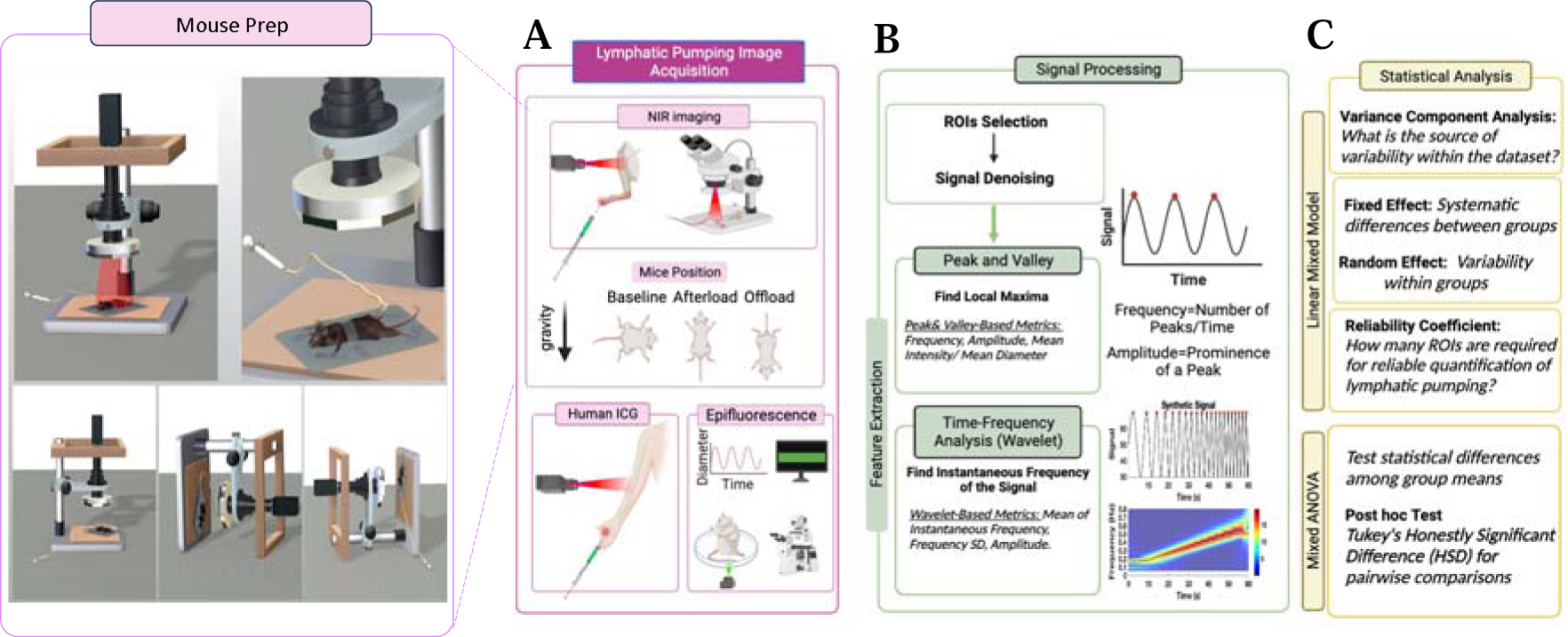
Image acquisition, signal processing and statistical analysis of real-time in vivo lymphatic pumping. (A) In mice, after footpad injection with NIR dye, NIRF lymphangiography is performed with the mouse and microscopy setup in three prone positions: baseline, afterload, and offload. In humans, after ICG dye injection in the interdigit webbed space, a NIRF handheld device is used to excite and image lymphatic vessels with the arm in a neutral position. In murine epifluorescence microscope lymphangiography, FITC in the hindlimb with skin removed is imaged in a single upright position. (B) Signals are processed in MATLAB by systematically selecting a row of ROIs, denoising, and creating a peak and valley plot. From the plot, classical amplitude and frequency are calculated. Time-frequency wavelet analysis is applied to the peak and valley plot, generating wavelet-based mean instantaneous frequency, frequency standard deviation, and amplitude. (C) For each type of experiment, a variance component analysis is performed. Based on this, experiment-informed fixed effects and random effects in a linear mixed model are determined, which is then used to determine statistical differences among group means with Tukey’s Honestly Significant Difference for posthoc pairwise comparisons. Decision theory is used to calculate a reliability coefficient for each metric in each experiment type to determine how many measurements per mouse or human are required for a reliable measurement.

To advance these methods, each ROI underwent wavelet analysis to determine the instantaneous contraction frequency over time (Figure 1B). Mean instantaneous frequency, frequency standard deviation, and amplitude are wavelet-based metrics that can be obtained from this analysis. For mouse NIRF and human ICG lymphangiography, we next used statistical methods to determine the significance and reliability of our measurements. For each method and measurement type, variance component analysis was performed to determine the sources of variability within the data (Supplemental Tables 1 and 2). Specifically, our statistical analysis revealed that the variability within individual mouse subjects accounts for 9-56% of the total variability the parameters of lymphatic function that we measured, while the variability within human subjects accounts for 25-84% of the total variance of our human data metrics. For almost all measurements, mouse-to-mouse differences are the primary source of variability in the data. This was larger than the variability attributable to age, sex, ROI, position, or interaction terms between these parameters (Supplemental Table 1). Exploration of these interactions helped determine the linear mixed model (see Methods; Statistical Analysis and Supplemental Table 1).

### Traditional peak-and-valley and wavelet time-frequency analyses reveal irregular patterns of lymphatic pumping in both mice and humans, while measurement reliability improves with the inclusion of more ROIs

NIRF imaging was performed in both mice and humans (Figure 2 A-B). In signal processing, ROIs were sequentially tiled along the length of the lymphatic vessel. Due to differences in tissue scattering and equipment resolution, 8 ROIs were used in recordings from mice and 4 were used in human recordings. Peak-and-valley plots of the imaging data reveal that fluorescence from the adult mouse has a continuous waveform that is both fairly regular and continuous, and has a mean frequency of 16.31±0.81 min^-1^ (Figure 2A,C). The wavelet analysis, which provides the instantaneous frequency of the signal, likewise shows a consistent instantaneous frequency with a mean of 18.18±0.78 min^-1^ for mice. On the other hand, the human peak-and-valley plot, while sometimes regular (Figure 2B), reveals frequencies that are an order of magnitude slower than the mouse (3.21±0.26 min^-1^ peak and valley and 1.91±0.2 min^-1^ wavelet-based mean of signal frequency). Unlike the continuous waveform of the mouse, the human waveform shows distinct peaks followed by a silent inter-peak interval (see peak/valley representative example, Figure 2B). This is reflected in the instantaneous frequency plot from the wavelet analysis, indicated by a region of non-zero frequency (Figure 2B) followed by a near-zero frequency between spikes.

**Figure 2:**
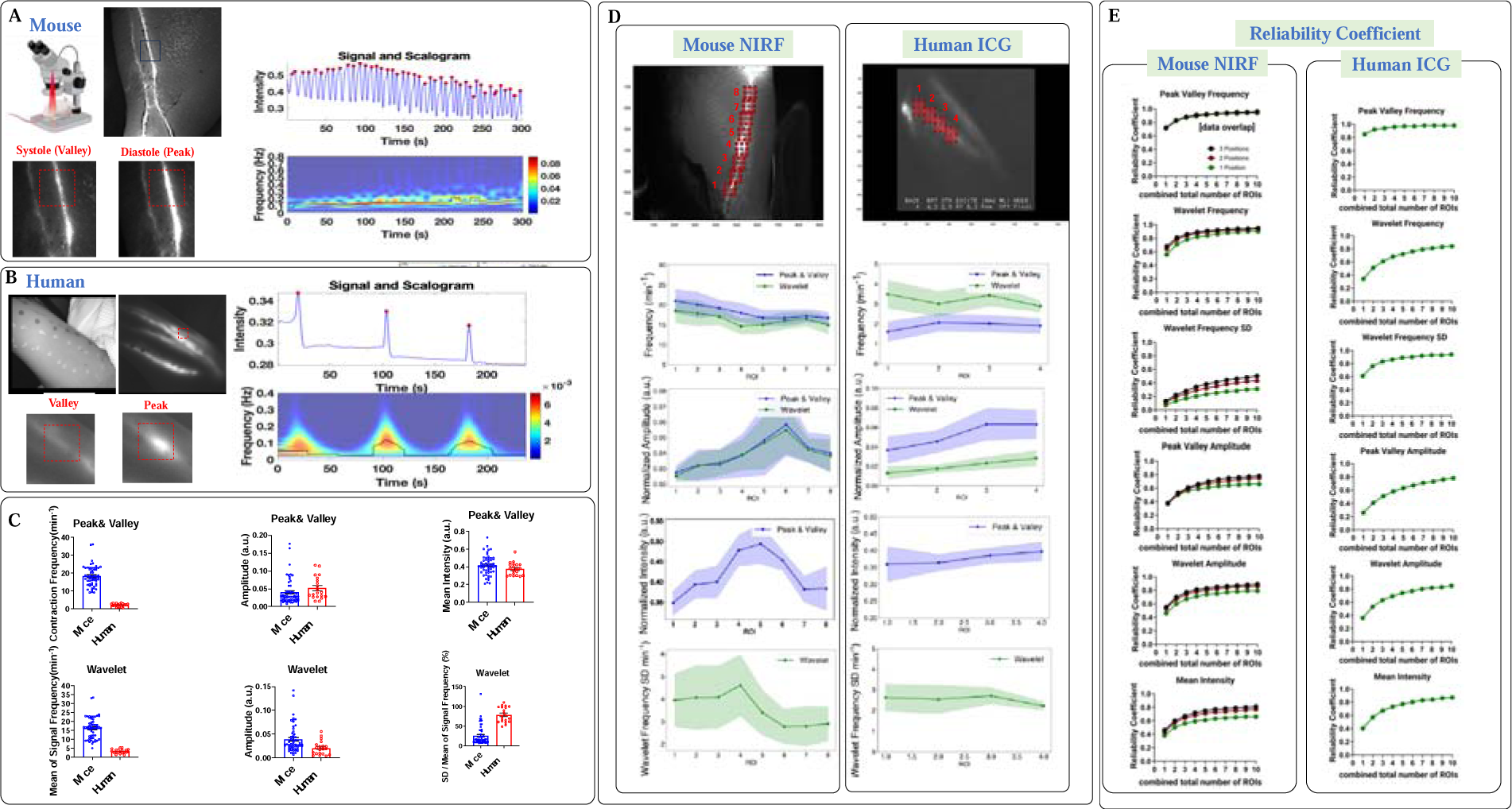
Near infrared lymphangiography measurement and analysis in mice and humans. (A) An example region of interest (red square) in the popliteal lymphatic network in systole and diastole. Peak and valley plot of the ROI showing the change in signal amplitude over time. Spectrogram of the peak and valley plot shows the instantaneous frequency over time. (B) An example region of interest (red square) in the forearm of a human patient showing the valley and the peak of ICG signal amplitude, with a corresponding peak and valley plot and spectrogram at that ROI. (C) The contraction frequency (from peak and valley analysis), mean intensity (from peak and valley analysis), mean instantaneous frequency (from wavelet analysis), standard deviation of instantaneous frequency (from wavelet analysis), and amplitude (from peak and valley and wavelet analysis) for mouse (left column with blue dots) and human (right column with red dots) lymphangiography (mean +/- SEM, N= 8 mice, 64 measurements, N = 5 humans, 20 measurements). Mouse data are plotted only for baseline position for comparison with human baseline position. (D) Measured parameters for mouse (left) and human (right data), for peak-and-valley (blue) and wavelet (green) methods showing moderate variation in some parameters by ROI. (E) The reliability coefficient for mean intensity, wavelet frequency, wavelet frequency SD, wavelet amplitude, peak and valley frequency, and peak and valley amplitude for mouse (left) and human (right) by number of ROIs measured for 1 (green line), 2 (red line), and 3 (black line) gravitational positions. As the values are derived from a model, we display reliability up to 10 ROIs, though 8 and 4 ROIs were originally measured from mice and humans, respectively.

While the frequency obtained by peak and valley versus wavelet analysis for the human data is one order of magnitude slower than for the mouse, and the peak and valley frequencies are consistent with wavelet analysis frequencies within each species (peak and valley 18 ± 0.78 min^-1^ and wavelet 16±0.81 min^-1^ for mouse; peak and valley 1.9 ±0.2 min^-1^ and wavelet 3.2±0.26 min^-1^ for human), the wavelet frequency standard deviation metric reveals much higher variations, capturing the irregular stop-start nature of the human lymphatic vessel contractility compared to mouse (frequency normalized SD (frequency SD min^-1^/ mean frequency min^-1^ * 100) human 78.6±4.3 vs mouse 25.3±3.2) (Figure 2C). Additional parameters captured in the peak and valley analysis include the mean intensity of the ROI, which is the normalized average intensity within the ROI (0.42±0.01 a.u. for mice vs. 0.38±0.02 a.u. for humans). The peak and valley amplitude, defined as peak prominence, is 0.04±0.01 a.u. for mice and 0.05±0.01 a.u. for humans. Similarly, the wavelet amplitude, quantifying the strength of the instantaneous signal, is 0.04±0.004 a.u. for mice and 0.02±0.003 a.u. for humans.

While Figures 2A and 2B illustrate the behavior of a single ROI, the lymphatic vessel is a chain of lymphangions and multiple ROIs can be tiled along the chain for analysis. There is variability in lymphatic contraction frequency, amplitude, signal intensity, and frequency irregularity between different ROIs in both mice and humans (Figure 2D), indicating that different ROIs may capture different contractile behavior and that the subjective choice of a single ROI along a chain of lymphangions may skew results or fail to capture part of the functional behavior. To investigate how many ROIs are required for reliable quantification of pumping metrics, we used decision theory to calculate the reliability coefficient for each type of measurement (Figure 2E). Increasing the number of measurements per mouse, both by increasing the number of ROIs and by increasing the number of gravitationally different positions, increases the reliability of the measurement (Figure 2E). For example, wavelet frequency calculated using only one ROI in one position has a reliability coefficient of only 0.56 and this increases up to 0.95 with eight ROIs and three body positions. This analysis also helped us identify at which point increasing the number of measurements per mouse has diminishing returns. For example, for peak and valley frequency, increasing the number of ROIs in a single position from one to five increases reliability from 0.71 to 0.91, while increasing the number of ROIs from five to 10 only increases the coefficient further to 0.94, a gain that may not be experimentally significant while increasing analysis time.

Each measurement type must be individually considered. For example, the measurement for wavelet frequency standard deviation has high variability and even with many ROIs and three positions, the reliability coefficient curve in mice does not saturate, predicting that additional measurements per mouse beyond those performed in this study would continue to increase the reliability of wavelet frequency standard deviation. For our human data, only one position was measured. Compared to the analogous measurement in mice, the human data have a lower reliability coefficient than the murine data for low numbers of ROIs. For example, the mouse reliability coefficient for wavelet frequency with four ROIs is 0.82, while the human reliability coefficient is 0.68. As seen in the murine data, increasing the number of ROIs increases reliability, with some metrics reaching saturation, while more variable measurements do not reach saturation even with 10 modeled ROIs. This suggests that the number of ROI measurements per human subject must be maximized to help improve the reliability of measurements and may be greater than the number needed in mice.

An important consideration is how to analyze repeat measurements in subjects when performing hypothesis testing between groups. One approach that has been used in the past is simply to use the mean of all individual data measurements of a single type when comparing groups, or to create a single data point for each measurement type in a subject by averaging the repeat measurements for that subject. In both approaches, valuable information is lost. Here, we employed a statistical approach that uses linear mixed models, in which repeat measurements in subjects are preserved as unique numbers and subject grouping can be achieved by using random intercepts by-subject in the model (see Methods; Statistical Analysis).

### Changes in position induces contractility variations between mice across different ages and sexes compared to baseline position

We next performed NIR imaging and analyses with mice in different positions with respect to gravity. For each video from a mouse in a single position undergoing NIRF imaging, eight ROIs were manually selected in MATLAB and tiled systematically in a row along the lymphatic vessel (Figure 3A). Measurements were then generated using both peak and valley and wavelet analysis. We analyzed the datasets several times with slight variations in the position of the eight tiled ROIs to test whether user input for ROI impacts results, and the dataset did not substantively change with slight variations in the tiling (data not shown). Linear mixed effects models including ROI, Sex, Age, Position, and some interaction terms between these (see Methods; Statistical Analysis) were used with Tukey’s HSD test for pairwise comparisons. In addition to the baseline position (mouse flat and prone; Figure 1A, 3A), we measured the mouse contractility with the head up (maximal afterload on the lymphatic vessel) and head down (maximal offload on the lymphatic vessel) (Figure 1A, 3A). In the afterload position, we hypothesized that the hindlimb lymphatic vessel would experience the greatest load as the vessel must pump against closed valves with the maximal column of fluid pressure above. In the offload position, the valves point in the same direction as the gravitational force. We hypothesized that the measured lymphatic vessel would experience the lowest load in this configuration. Surprisingly in the adult mouse (Figure 3B), there was no difference in contraction frequency between the afterload position (20.3±3.8 min^-1^ for peak-and-valley and 20.0±3.2 min^-1^ for wavelet) and baseline measurements (19.2±3.8 min^-1^ for peak-and-valley and 17.3±3.2 min^-1^ for wavelet). Compared to baseline prone position, the amplitude of pumping and mean intensity of the signal in the afterload position is lower in adult mice (Figure 3B), consistent with the vessel being able to move less fluorescent dye in these increased load conditions. We did not see a recovery of amplitude or signal intensity when mice were returned from the afterload to the offload position, though the frequency did decrease.

**Figure 3:**
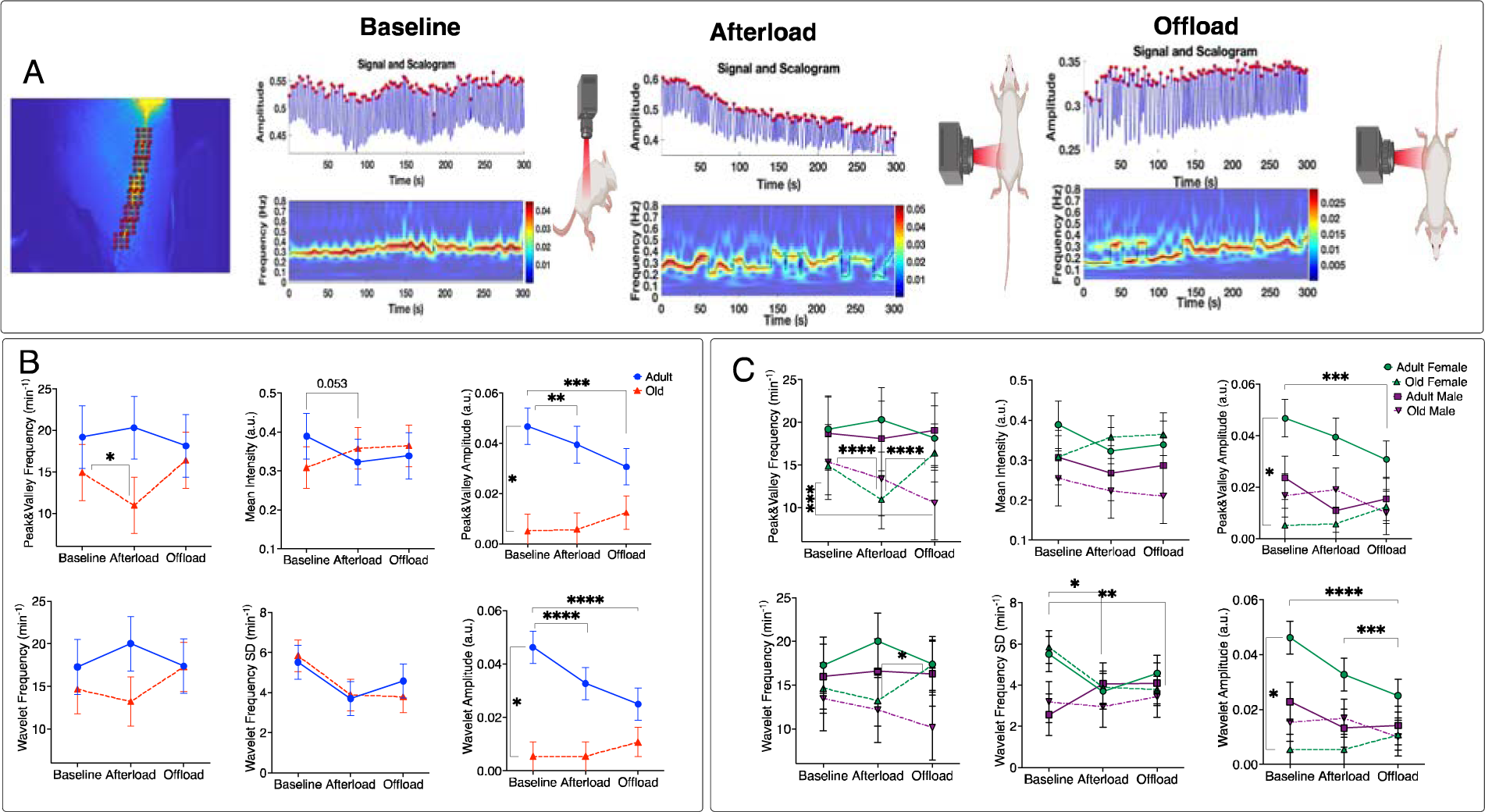
Near Infrared Lymphangiography Measurement in Mice Across Age, Sex, and Lymphatic Position. (A) An example image from a series of pictures comprising the 5-minute lymphangiography recording showing 8 regions of interest tiled across the lymphatic vessel. The peak-and-valley plot and spectrogram are shown in the baseline position, afterload position (head up), and offload position (head down). (B) The model estimate +/- SEM from linear mixed modeling for peak and valley frequency (min^-1^), wavelet frequency (min^-1^), peak and valley amplitude (a.u.), wavelet amplitude (a.u.) and peak and valley mean intensity (a.u.) are given for the baseline, afterload and offload positions in adult (blue line) and old (red line) mice (N= 8 adult mice, N = 8 old mice), data collapsed across sex. (C) The same data as in (B) is shown split by sex, with green solid line adult female (N = 5), green dotted line old female (N = 4), purple solid line adult male (N = 3), purple dotted line old male (N= 4).

The behavior of old mice is different than adult mice (Figure 3B), with overall frequencies and amplitudes lower in all three positions for old mice. While baseline position measurements are different between age groups, the stress of the afterload position reveals an even larger difference between the groups. When faced with afterload, the contraction frequency decreases in old mice (old mouse afterload to old mouse baseline contrast for peak and valley frequency min^-1^: mean difference =-2.95, SE =0.77, p =0.002). This demonstrates the value of measuring the behavior of the dynamic physiological system of lymphatic vessels under multiple load conditions to explore the full gamut of system behavior. The decrease in lymphatic contraction frequency in old mice under afterload conditions could be interpreted as pump failure due to the additional load (i.e., the system is unable to compensate for the increased load by increasing pumping strength). We further analyzed our data by sex, which revealed additional differences between sexes. While in Figure 3B, in both peak and valley and wavelet frequency graphs, the old mice (red line) show a pattern of decreased pumping frequency with afterload position compared to baseline position, with a recovery increase in the offload position (not all of these comparisons reach significance in difference in Figure 3B), it is apparent in Figure 3C that the lymphatic pumping behavior splits by sex in the old mice. Notably, the recovery of peak and valley frequency in the offload position in old mice is only in female mice (green dotted line) (p <0.001). When male and female mice are analyzed together, the recovery of frequency in offload is not significant. The old male mice (dotted purple line in Figure 3C) continue to decrease in frequency in the offload position and there is a significant decrease in peak and valley frequency between old male mice in the baseline and offload positions (peak and valley frequency min^-1^ baseline to offload: mean difference =4.79, SE =1.2, p =0.0053).

### Wavelet analysis of murine intravital epifluorescence microscopy reveals differences in lymphatic pumping regularity not revealed by traditional analysis methods

To test whether our wavelet analysis can be applied to lymphatic contractility measurements from intravital epifluorescence microscopy, we re-analyzed previously published intravital epifluorescence microscopy data in young and old mice[33] that was previously quantified using only peak and valley analysis (Figure 4A). Both traditional peak-and-valley and wavelet analyses suggest there is no statistically significant difference in frequency between young and old mice (e.g., 8.3 ± 0.9 for young and 7.7 ± 0.9 for aged mice, based on peak-and-valley analysis). The value of frequency obtained by peak and valley analysis is also dependent on a user-selected threshold of peak change, in this re-analysis 2 µm of vessel diameter. Both peak and valley techniques and wavelet analysis show that the amplitude of contractions significantly decreases with aging (13.9 ± 1.7 µm in young and 4.5 ± 1.1 µm in old for peak and valley; and 10.4±1.6 µm in young and 3.6±0.8 µm in old for wavelet, p < 0.05, unpaired t-test).

**Figure 4:**
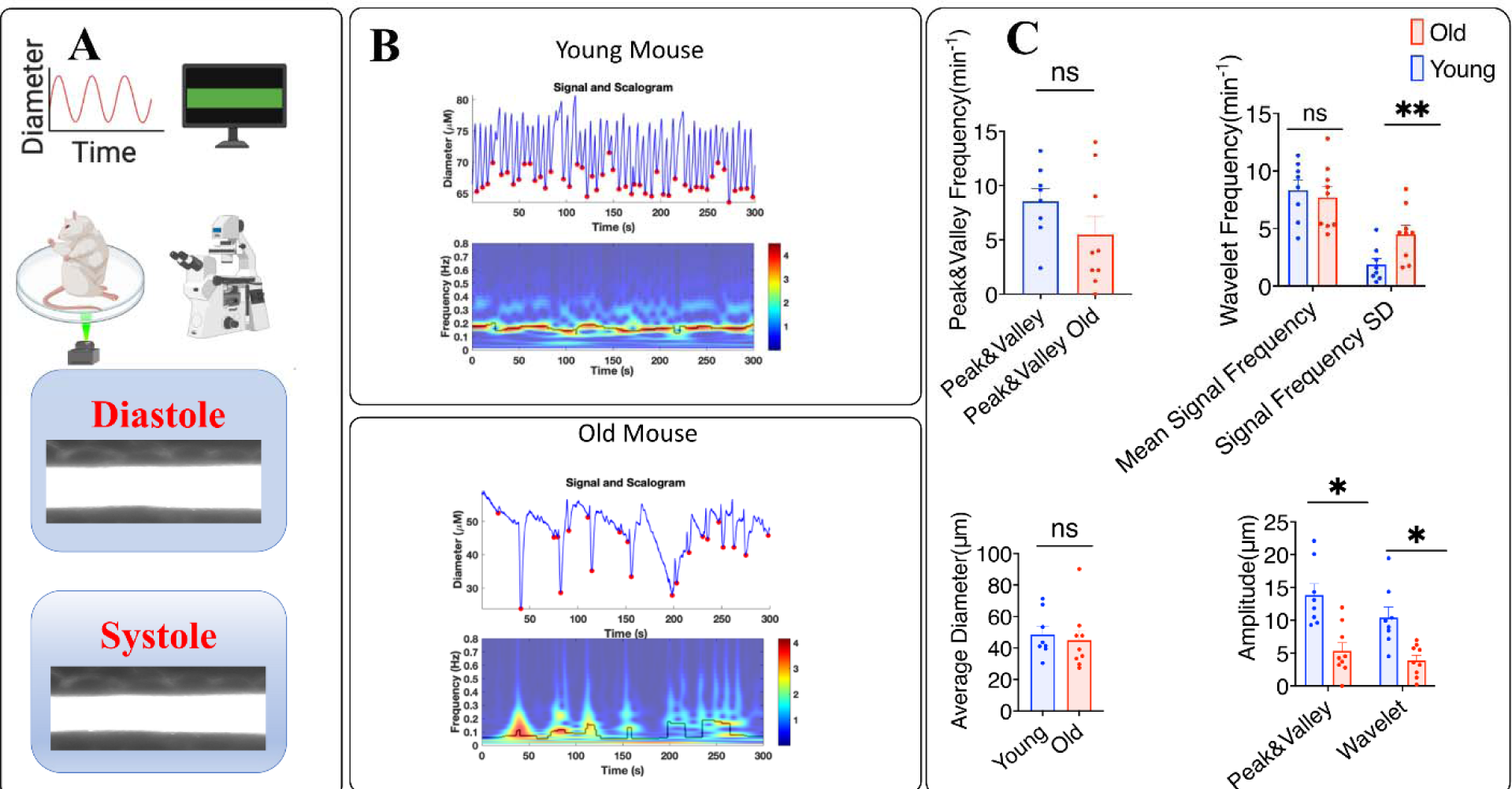
Intravital epifluorescence microscopy. (A) A lymphatic vessel afferent to the popliteal lymph node in a live mouse is filled with fluorescein isothiocyanate dye by interdermal injection of the mouse foot and imaged on an epifluorescence microscope. A single region of interest is captured and diameter changes over time are recorded. An example lymphatic vessel in diastole and systole is shown. (B) Example data in a young mouse. The average vessel diameter over time is measured over 300 seconds. The spectrogram below shows the instantaneous frequency throughout the recording. Below, example diameter vs. time and instantaneous frequency vs. time for an old mouse. (C) Mean +/- SEM of contraction frequency (peak and valley analysis), frequency (peak and valley analysis), amplitude (peak and valley and wavelet analysis), and average diameter (peak and valley analysis) for young (blue), N = 9 mice and old (red), N = 8 mice.

While the peak and valley plot visually looks irregular in the old mice, the peak and valley plot is smooth and regular in young mice (Figure 4B). Previous methods for quantifying lymphatic contractility fail to capture this change to more irregular contraction behavior with age. Interestingly, the wavelet frequency standard deviation, which measures the deviation from the mean of the instantaneous frequency, is significantly higher in the old mouse group due to large changes in the instantaneous frequency of lymphatic pumping (1.9 ± 0.5 min^-1^ for young vs 4.8 ± 1.0 min^-1^ old group, p < 0.01) (Figure 4C). This novel parameter can capture irregularities such as the start-stop nature of lymphatic contraction in aged mice. In this re-analysis of prior data, only one ROI per mouse was available, so we did not explore reliability analysis with multiple repeat measures. In intravital epifluorescence microscopy, time frequency analysis using wavelets is a powerful tool that can be especially useful in the analysis of aged or diseased mice to capture changes in the regularity of lymphatic pumping.

## DISCUSSION

The past two decades have witnessed advancements in non-invasive imaging modalities for imaging lymphatic vessels. Among these techniques, NIRF lymphatic imaging and ICG lymphography, which are closely related techniques, have gained popularity for applications in small animal models of lymphatic disease and the clinical evaluation of human lymphatic dysfunction, respectively[17, 22, 24, 25, 28, 34]. Typically, the analysis of NIRF signals is performed by selecting an ROI and analyzing the average signal intensity over time, which is associated with the intrinsic contraction of the lymphatic vessels. Functional metrics based on peak and valley analysis, such as frequency, ejection fraction, emptying rate, pumping pressure and fractional pump flow, have been used to analyze NIRF signals in lymphatic diseases[35, 36]. While these metrics offer valuable insights, the peak and valley-based metrics lack temporal resolution.

Some recent studies show the benefit of more advanced signal processing techniques for understanding lymphatic contractility. For example, wavelet analysis has shown promise for demonstrating the entrainment of lymphatic contraction in an isolated vessel experimental setup with applied cyclic fluid pressure[37]. Wavelet analysis is commonly used for electroencephalograms (EEGs)[38], and has also been used to analyze and extract features of complex biomedical signals such as electrocardiograms (ECGs)[39, 40] and electromyograms (EMGs)[41, 42]. Unlike Fourier transforms, which are suitable for temporally stationary signals, wavelet transforms—with their varying frequency and length—are better suited for signals with time-varying components, thus providing multi-resolution characteristics that provide insights into both temporal and frequency aspects of the signal. In particular, wavelet analysis can be useful when there is irregularity in lymphatic contractility. The novel contractility parameter of instantaneous frequency SD can capture the irregularity of lymphatic contractility, which is missed in traditional peak-and-valley analysis.

In NIRF imaging studies of the human lymphatic system, the parameters of lymphatic propulsion frequency and velocity are valuable and innovative physiological metrics that have been used for assessing lymph flow[43, 44]. However, the calculation of these parameters necessitates the measurement of the distance traveled by fluorescent markers. The data captured in NIRF imaging is in the form of 2-dimensional images, but the fluorescence originates from a 3-dimensional lymphatic vessel that varies in position along the Z-plane of the human tissue. Consequently, the 2-dimensional distance derived from projecting photons from all Z-planes in the collected data overlooks changes in the tissue depth of the imaged vessel. This omission restricts the reliability of velocity comparisons between individuals due to the unaccounted variance in vessel depth. Therefore, to ensure a more consistent and comparative analysis across different subjects, we prioritize the frequency and amplitude of fluorescence as our principal derived parameters, as they offer a more straightforward comparison between subjects without the complexities introduced by depth variations.

While most studies have used a single or few ROIs when analyzing lymphatic contractility via NIRF imaging, our statistical analysis using a linear mixed model suggests that such quantification may not be reliable. Reliability Coefficient testing assigns a value between 0 to 1 to a given test, with R= 1.00 indicating perfect reliability[45]. Test-retest reliability in animal research has great variability, with certain measurements notoriously unreliable between different experimenters and groups, such as in behavioral studies[46, 47]. For clinical measurements used in patient care, a reliability coefficient above 0.9 has been proposed as acceptable[48]. To achieve reliable measurements for most of our experimental parameters in mice, we need at least five measurements from different ROIs, while more variable metrics require an even greater number of ROIs. The measurement reliability is further increased by studying lymphatic vessels in multiple positions with respect to the force of gravity.

The use of multiple ROIs is advantageous as it allows the analysis of lymphatic pumping at different positions along the chain of lymphangions. For our human lymphangiography measurements, the highest number of measurements that were available (4 ROIs) did not meet clinical reliability standards (reliability coefficient > 0.9) for our functional metrics, demonstrating that development of a reliable bedside lymphangiography test requires more measurements per subject than is typically performed for ICG lymphangiography anatomic studies. We showed that decision theory can be used to find the optimal number of measurements per subject needed to create a reliable measurement for each experimental parameter, and that this number differs between mice and humans, even when using a similar NIRF lymphangiography technique. It is notable that in both mice and humans, lymphatic contractility greatly varies between subjects and subject variability is a dominant factor in the total attributable measurement variability of our experiments. This highlights the need for making measurements within a single subject as reliable as possible. We showed that increasing the number of ROIs measured and handling the repeated measurements thoughtfully with a mixed model is a way to increase reliability. Increasing reliability is key in the development of tests that can be safely implemented in clinical decision-making. We also found that variance component analysis can be used to identify which parameters in the data (for example, subject number) contributes to the greatest source of variability in the data. Variance component analysis then informs an optimal linear mixed model for statistical analysis between different groups.

Our data also show differences in lymphatic contractility parameters between mice and humans. The overall frequency of lymphatic contraction in the human data is lower than that of adult mice, while the irregularity of contractility as measured by normalized frequency SD is greater. There is emerging evidence that anesthetics change lymphatic contractility[49], and notably, mice were under anesthesia during lymphangiography while humans were not. While the large differences in lymphatic function between mice and humans are most likely driven by species differences, anesthetic factors are worth bearing in mind and testing in greater detail in the future.

The lymphatic system needs to adjust to different fluid loading scenarios. Gravity is one important factor that could alter fluid dynamics in the interstitial space and vessels, creating different hydrostatic pressure gradients (e.g., in standing versus supine positions). In clinical settings, it is common to elevate the lower part of a patient’s body above their head in some surgical procedures, which exposes lower extremity lymphatic vessels to favorable hydrostatic pressure gradients and head-and-neck lymphatics to unfavorable pressure gradients. Experiments simulating the effects of microgravity on lymphatic contractility, along with computational modeling, also suggest that gravity alters lymphatic contractility[50, 51]. In the present study, we utilized NIRF imaging and designed a specialized microscope fixture to investigate the effects of gravity on lymphatic contractility. Employing a quantitative approach, we assessed the differences in lymphatic function between older and younger mice when placed in various positions relative to gravity. Our analysis indicates that exposing adult mice to a head-up position, which maximizes the effect of gravity on the lower limbs, had limited effect on lymphatic contractility. Similarly, placing adult mice head-down, where gravity facilitates flow by inducing a favorable hydrostatic pressure, does not significantly alter contraction frequency. This is likely due to the small size of mice, which limits the magnitude of the gravitational forces. However, older mice exhibit decreased contractility in the head-up position compared to their baseline, revealing age-related differences in response to gravitational changes. We suggest incorporating measurements in different gravitational positions to standard NIRF mouse lymphangiography to more fully capture the range of behavior of lymphatic contractility. Furthermore, splitting the data by individual sex shows differences in some metrics between groups, suggesting that sex is an important biological demographic variable in lymphatic contractility quantification and should not be ignored.

In lymphatic measurements, a persistent challenge and objective is to minimize animal-to-animal variability[52] and handle regional heterogeneity[53]. NIRF lymphangiography in mice has been criticized for its unreliability and high variability compared to *ex-vivo* lymphatic measurement methods[54]. We show that incorporating more measurements per animal, in the form of more ROIs and measurements in multiple positions can generate highly reliable data with test-retest coefficients of experimental parameters >0.9, considered adequate even for clinical tests. In the emerging field of human ICG lymphangiography, increasing measurements per human subject may be what is needed to evaluate ICG lymphangiography as a reliable clinical tool that could be utilized in clinical practice to quantify lymphatic function in at-risk patients and those already with the burden of lymphatic disease. Finally, wavelet transform analysis is a powerful method to reveal irregularity in lymphatic contraction that is not revealed by traditional lymphangiography methods, which is especially crucial in the study of human subjects, and mice with disease phenotypes. Combined, implementing these new methods can bring human and murine NIRF lymphangiography to the forefront as a more reliable tool in testing lymphatic function in patients and preclinical models.

## METHODS

### Mouse Model

C57BL/6 mice, female and male, 6 months old, were obtained from the Cox-7 animal facility operated by the Edwin L. Steele Laboratories, Department of Radiation Oncology at the Massachusetts General Hospital (MGH). C57BL/6 mice, female and male, 18 months old, were obtained from the National Institute on Aging. Both ages of mice were housed at the MGH Center for Comparative Medicine facilities, Charlestown Navy Yard. Animal protocols were approved by the Institutional Animal Care and Use Committees (IACUC) at MGH, and all facilities are accredited by the Association for Assessment and Accreditation of Laboratory Animal Care International (AAALAC).

#### Imaging Lymphatic Pumping

A combination of ketamine and xylazine was used to anesthetize mice. The fur was removed using a depilatory cream, and 5μL of Dextran (10K), Flamma® 774 (BioActs) was injected intradermally into the dorsal aspect of the mouse paw. NIRF imaging, as previously described[55], was used to image the saphenous lymphatic vessels afferent to the popliteal lymph node. Briefly, the imaging system is equipped with a ×6.5 Zoom lens (Navitar), a Prosilica GT2750 camera (Allied Vision Technology), and an ICG-B emission filter (832/37, Semrock). A 760 nm laser-emitting diode (Marubeni), filtered through a 775/50 bandpass filter (Chroma), was used to excite the dye. To study how gravity affects lymphatic pumping, a custom-designed microscope fixture was integrated with the imaging system. This setup allowed the positioning of mice to experience different gravitational forces during imaging. The recordings were conducted in five-minute intervals, during which the mice were positioned in three distinct postures: 1) prone, 2) head-up (their heads elevated above their bodies), 3) head-down (their heads positioned lower than their bodies).

#### Human ICG lymphography

Videos were obtained from Dr. Dhruv Singhal at BIMDC as previously described. Briefly, 0.1 mL stock solution (2.5 mg/mL) ICG solution (Akorn) was sterilely mixed with 2.5 mg albumin and injected intradermally. Multiple injections were introduced as previously described[23]. An injection overlying the cephalic vein was introduced to image the upper arm. Only upper arm images were used in this study. A NIRF imager (Hamamatsu PDE Neo II, Mitaka USA) was used to visualize superficial lymphatic vessels with the ‘mapping’ mode of the device. Institutional review board approval was obtained (2021P000209) for this prospective lymphangiography analysis. Consent was obtained from every patient to perform the studies.

#### Signal Processing

A custom-written MATLAB code was used to analyze images recorded at 10 fps. The ROIs, consisting of a 50×50 pixel area (mouse) or 100 x 100 pixel area (human), were selected along the lymphatic vessels. The normalized average intensity (0 < Intensity < 1) within each ROI was recorded at each time point. To denoise the signal, we used a wavelet denoising technique based on the Block James-Stein method, which yields optimal global and local adaptivity[56]. The denoised signal was analyzed based on two approaches: 1) Peak and Valley: Frequency was detected based on the local maxima in the signal, and amplitude was determined based on peak prominence, which measures how the peak’s height stands relative to other peaks. 2) Multiresolution Time-Frequency Analysis: The instantaneous frequency of the signal was detected through the wavelet transformation of the signal, identifying the wavelet component with the highest magnitude at each time point. The instantaneous amplitude of the signal was also correlated with the magnitude of the corresponding wavelet component with the highest magnitude. Specifically, continuous wavelet transformation employing the Morse mother wavelet was utilized, which is well-suited for analyzing non-stationary signals that exhibit frequency variations over time. The average and standard deviation of frequency and amplitude in the instantaneous frequency and amplitude of the signal were used as metrics for lymphatic pumping.

## DETAILED PROCEDURES

### Mouse preparation

1. Place the mouse on a scale to weigh it. Anesthetize the mouse using the ketamine-xylazine mixture (100 mg/10 mg per kg body weight) injected subcutaneously in the mouse’s lower left quadrant.

- ***Note:*** *Maintain the body temperature at 37°C throughout the procedure using a heating pad set to 39°C*.
2. Once the mouse is in an adequate plane of anesthesia, apply ophthalmic ointment to both eyes, then shave the mouse’s right hindlimb toward the thigh initially and then in the opposite direction, toward the footpad, for optimal fur clearance.
3. Remove any remaining hair from the right lower hindlimb by using depilatory cream. Remove the cream after 60 s with sterile filter-sterilized saline, and Kim wipe to prevent chemical skin burns.
4. Inject 5μL of Dextran (10K) Flamma® 774 (BioActs) intradermally into the dorsal aspect of the mouse paw, between the 2^nd^ and 3^rd^ digits.

- ***Note:*** *Injections on the ventral aspect of the footpad are usually subcutaneous rather than intradermal, take longer to fill the lymphatic network, and are thus not preferred*.

### Mouse staging

5. Place the mouse on a rectangular-shaped light-controlled aluminum sheet in the prone position and tape down all extremities and the tail with the right hindlimb at a 45-degree angle away from the body— place tape along the upper back to secure the mouse for positional changes.
6. Tape the light-controlled aluminum sheet onto the warming pad with dual-sided mounting tape (Gorilla) and frame the edges with tape to ensure the sheet is pressed firmly against the heating pad.

### Imaging

7. The afferent lymphatic vessels to the popliteal lymph node are imaged using near-infrared imaging using a previously described NIRF setup.
8. The imaging system is composed of a ×6.5 Zoom lens (Navitar) and a Prosilica GT2750 camera (Allied Vision Technology) with an ICG-B emission filter (832/37, Semrock).
9. A custom-built ring light source is used to excite the dye using 760 nm high-power laser-emitting diodes (Marubeni) filtered through a 775/50 bandpass filter (Chroma).
10. A custom-written MATLAB code is used to analyze images. The code is available upon request from the authors.

### Positional Maneuvers

11. Take a 5-minute recording of the mouse in the Baseline position (imaging system flat).
12. Move the apparatus using the supportive wooden frame so the mouse is in the afterload position and allow 2 minutes for equilibration, then take a second 5-minute recording.
13. Move the apparatus again to place the mouse in the offload position, allow 2 minutes for equilibration, and then take a third 5-minute recording.

### Statistical Analysis

#### Variance Components Analysis and Generalizability Theory

The primary aims of this study were to evaluate the percent of variance attributable to each measurement facet (mouse, position, ROI, age, sex) for each outcome (wavelet frequency and standard deviation, wavelet amplitude, mean intensity, peak distance, and peak valley frequency and amplitude) and to assess the reliability of the measurements. The attributable variance (expressed as a percentage) for each measurement facet was assessed for each outcome using a linear mixed effects model that included each facet as its own random intercept, as well as random intercepts for all two-way interactions between facets (Initial Model in Supplementary Table 1). Within-mouse two-way interactions that were not crossed (e.g., mouse:sex and mouse:age) were excluded as they could not be estimated because each mouse was only one sex and one age.

To assess measurement reliability, the full model from above was adjusted for each outcome to remove any within-mouse interaction terms that had zero or near-zero variance (Refined Model in Supplementary Table 1). Using the mouse identifying number (Mouse_ID) as the object of measurement, the simplified model for each outcome was input into the “gstudy” and “dstudy” functions of the “gtheory” package in RStudio. Generalizability coefficients were calculated using relative error variances, a measurement of the “noise” around a data point, and components from the “gstudy” output[57]. The generalizability coefficients were used to assess reliability as a function of the number of ROIs and position measurements.

#### Hypothesis Testing

The secondary aim of this study was to examine associations between mouse characteristics and the near-infrared imaging outcomes. Descriptive statistics for continuously scaled variables are reported as mean ± standard error of the mean. These data were analyzed using a linear mixed effects model, with a random intercept by mouse to account for the repeated nature of data collection. Models also included a two-way multiplicative interaction between ROI and sex, and a three-way interaction between age, sex, and position. The following structure in R was used:

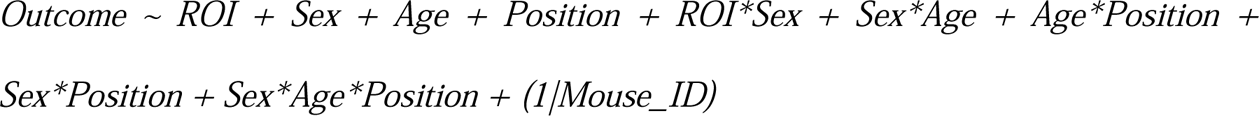

Model estimates are reported as predicted mean ± standard error. Tukey’s HSD test was used to assess pairwise comparisons of the three-way interaction between age, sex, and position, and the two-way interaction between age and position. Results were evaluated using two-sided p-values, with values less than 0.05 considered statistically significant.

Power calculations were conducted to illustrate the sample size necessary to detect a statistically significant difference between age groups (adult vs. old) for each outcome for reliability = 0.90 and reliability = 0.70. These values were chosen to represent high and low levels of reliability in outcome measurements. Low levels of reliability obscure the differences between groups causing a reduction (i.e., attenuation) of observeable effect size difference. To obtain the expected effect sizes that would be observed under these reliability conditions, the observed effect sizes and reliabilities from each model were disattenuated and attenuated to generate theoretical effect sizes. In this way, the effect sizes that would be observed under different levels of reliability could be calculated. To estimate power, a two-sample independent t-test was used to determine the n per group needed to achieve 0.80 power with a type I error rate of 0.05.

## Author Contributions

Mohammad S. Razavi, Ph.D Experimental design, data collection, data figures, data analysis, manuscript writing, manuscript revision

Katarina J. Ruscic M.D., Ph.D. Experimental design, data collection, data figures, data analysis, manuscript writing, manuscript revision

Elizabeth G. Korn, M.P.H. Statistical analysis, manuscript revision

Marla Marquez, B.A. Data collection, data figures, manuscript revision

Timothy T. Houle, Ph.D. Statistical design, statistical analysis, manuscript revision

Dhruv Singhal, M.D. Data collection, manuscript revision

Lance L. Munn, Ph.D. Experimental design, figure design, manuscript revision

Timothy P. Padera, Ph.D. Experimental design, manuscript revision

## Code availability

The code is available upon request from the authors.

## Acknowledgements

This works was supported by NIH grants F32HL156654 (MSR), R21AG072205 (TPP), R01HL128168 (LLM,TPP), R01CA284372 (TPP), R01CA284603 (LLM, TPP), R21EB031982 (LLM) and R01CA247441 (LLM) as well as support from the Rullo Family MGH Research Scholar Award from the MGH Research Institute (TPP). This work was also supported by the National Institutes of Health via Harvard Anesthesia T32-GM007592, by a grant from the International Anesthesia Research Society awarded to KJR, and by the Harvard Medical School Eleanor and Miles Shore Award to KJR.

---

**Supplemental Table 1:**
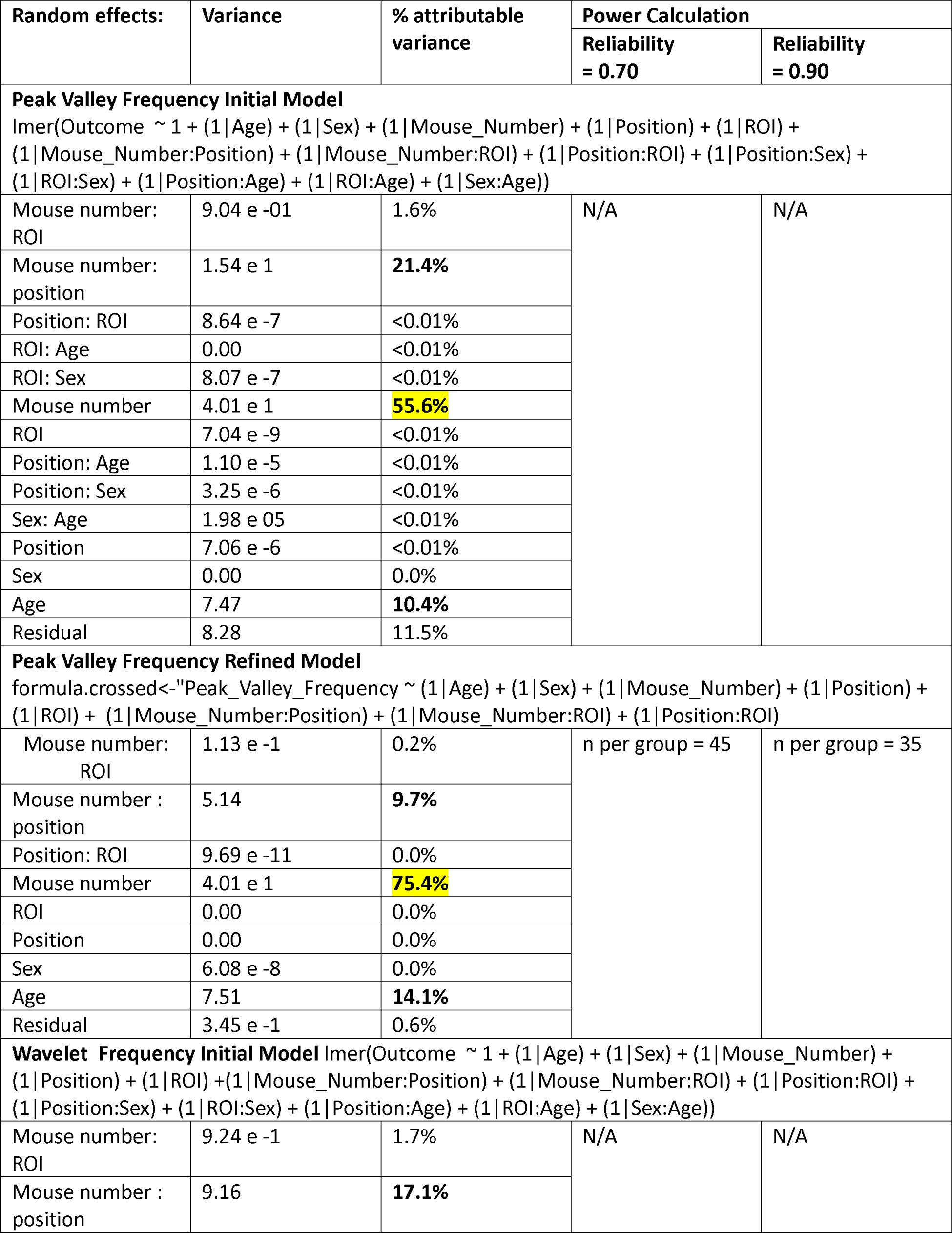
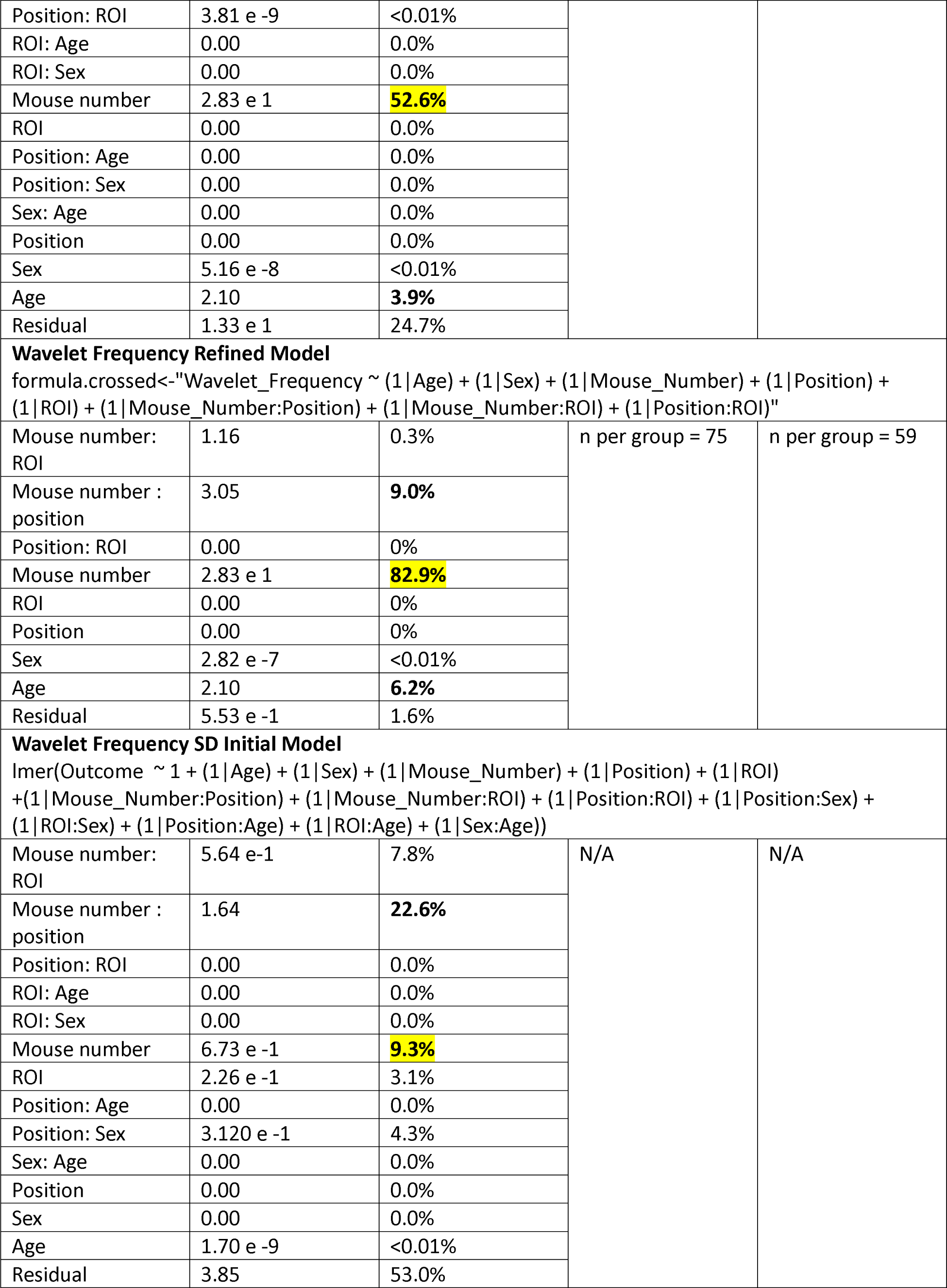
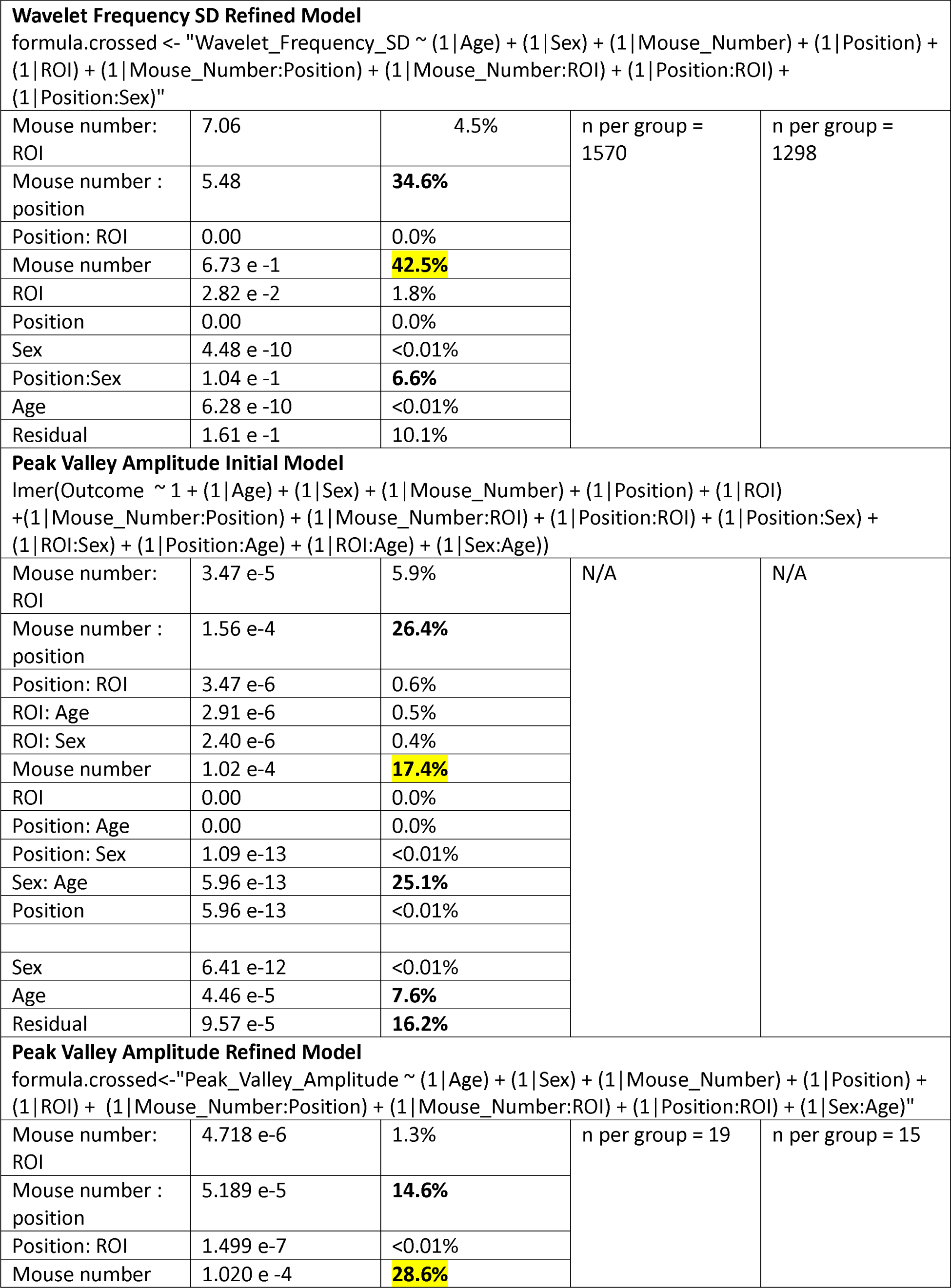
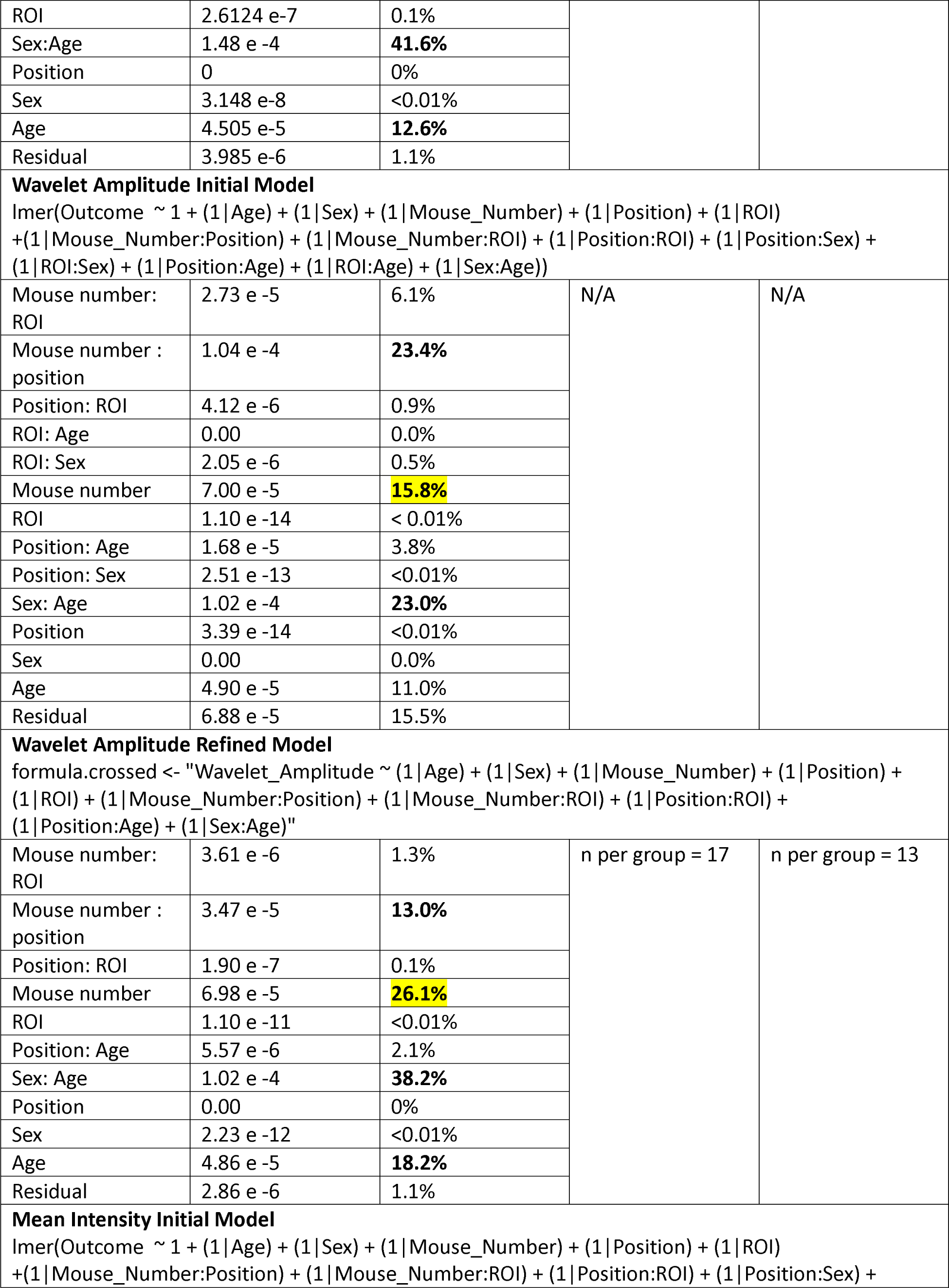
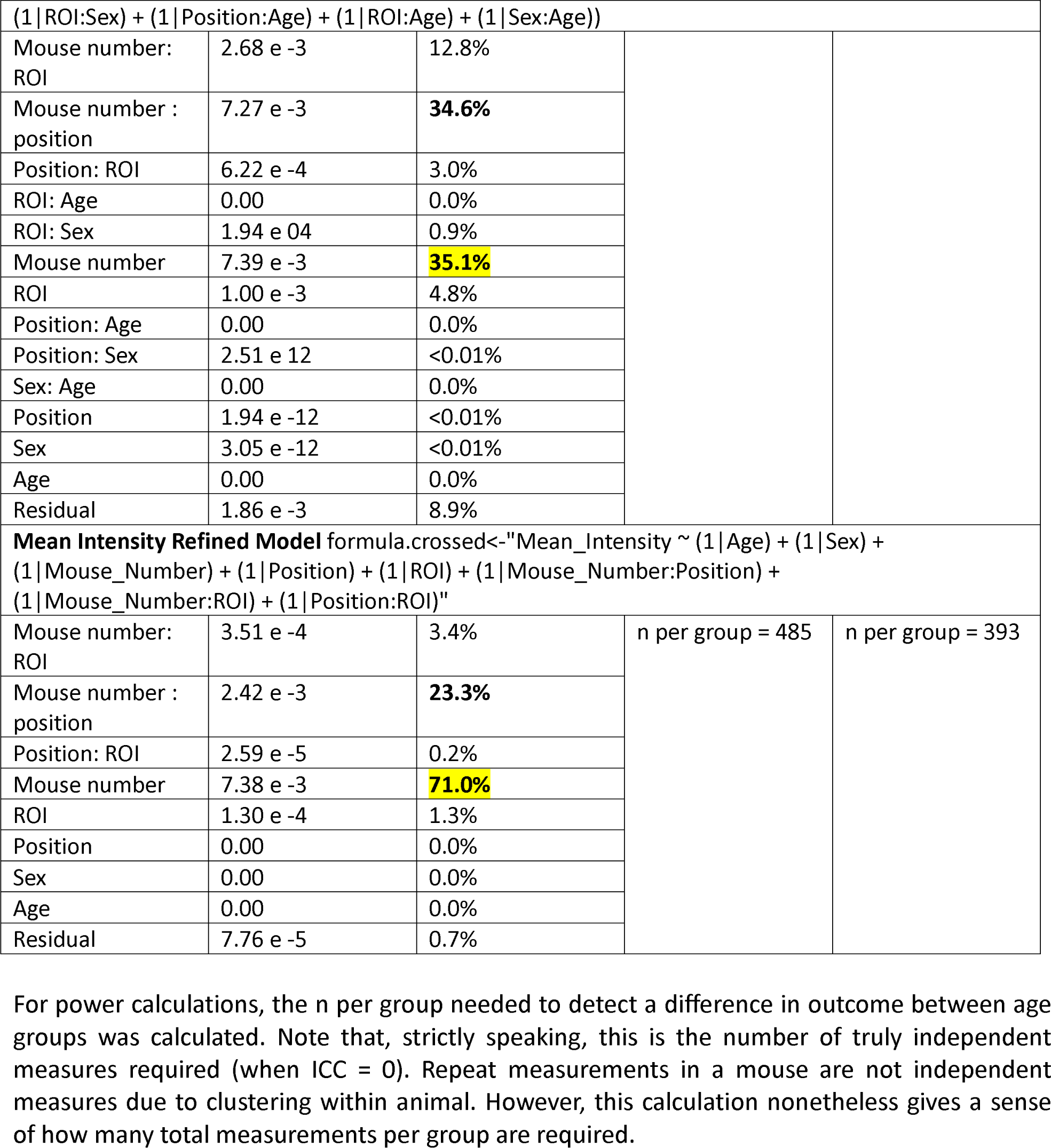
Variance Components Analysis: Murine Data, Initial and Refined Models.

| Random effects: | Variance | % attributable variance | Power Calculation |  |
| --- | --- | --- | --- | --- |
|  |  |  | Reliability = 0.70 | Reliability = 0.90 |
| Peak Valley Frequency Initial Model |  |  |  |  |
| lmer(Outcome ~ 1 + (1 \| Age) + (1 \| Sex) + (1 \| Mouse_Number) + (1 \| Position) + (1 \| ROI) + (1 \| Mouse_Number:Position) + (1 \| Mouse_Number:ROI) + (1 \| Position:ROI) + (1 \| Position:Sex) + (1 \| ROI:Sex) + (1 \| Position:Age) + (1 \| ROI:Age) + (1 \| Sex:Age)) |  |  |  |  |
| Mouse number: ROI | 9.04 e -01 | 1.6% | N/A | N/A |
| Mouse number: position | 1.54 e 1 | 21.4% |  |  |
| Position: ROI | 8.64 e -7 | <0.01% |  |  |
| ROI: Age | 0.00 | <0.01% |  |  |
| ROI: Sex | 8.07 e -7 | <0.01% |  |  |
| Mouse number | 4.01 e 1 | 55.6% |  |  |
| ROI | 7.04 e -9 | <0.01% |  |  |
| Position: Age | 1.10 e -5 | <0.01% |  |  |
| Position: Sex | 3.25 e -6 | <0.01% |  |  |
| Sex: Age | 1.98 e 05 | <0.01% |  |  |
| Position | 7.06 e -6 | <0.01% |  |  |
| Sex | 0.00 | 0.0% |  |  |
| Age | 7.47 | 10.4% |  |  |
| Residual | 8.28 | 11.5% |  |  |
| Peak Valley Frequency Refined Model |  |  |  |  |
| formula.crossed<-"Peak_Valley_Frequency ~ (1 \| Age) + (1 \| Sex) + (1 \| Mouse_Number) + (1 \| Position) + (1 \| ROI) + (1 \| Mouse_Number:Position) + (1 \| Mouse_Number:ROI) + (1 \| Position:ROI) |  |  |  |  |
| Mouse number: ROI | 1.13 e -1 | 0.2% | n per group = 45 | n per group = 35 |
| Mouse number : position | 5.14 | 9.7% |  |  |
| Position: ROI | 9.69 e -11 | 0.0% |  |  |
| Mouse number | 4.01 e 1 | 75.4% |  |  |
| ROI | 0.00 | 0.0% |  |  |
| Position | 0.00 | 0.0% |  |  |
| Sex | 6.08 e -8 | 0.0% |  |  |
| Age | 7.51 | 14.1% |  |  |
| Residual | 3.45 e -1 | 0.6% |  |  |
| Wavelet Frequency Initial Model |  |  |  |  |
| lmer(Outcome ~ 1 + (1 \| Age) + (1 \| Sex) + (1 \| Mouse_Number) + (1 \| Position) + (1 \| ROI) +(1 \| Mouse_Number:Position) + (1 \| Mouse_Number:ROI) + (1 \| Position:ROI) + (1 \| Position:Sex) + (1 \| ROI:Sex) + (1 \| Position:Age) + (1 \| ROI:Age) + (1 \| Sex:Age)) |  |  |  |  |
| Mouse number: ROI | 9.24 e -1 | 1.7% | N/A | N/A |
| Mouse number : position | 9.16 | 17.1% |  |  |
| Position: ROI | 3.81 e -9 | <0.01% |  |  |
| ROI: Age | 0.00 | 0.0% |  |  |
| ROI: Sex | 0.00 | 0.0% |  |  |
| Mouse number | 2.83 e 1 | 52.6% |  |  |
| ROI | 0.00 | 0.0% |  |  |
| Position: Age | 0.00 | 0.0% |  |  |
| Position: Sex | 0.00 | 0.0% |  |  |
| Sex: Age | 0.00 | 0.0% |  |  |
| Position | 0.00 | 0.0% |  |  |
| Sex | 5.16 e -8 | <0.01% |  |  |
| Age | 2.10 | 3.9% |  |  |
| Residual | 1.33 e 1 | 24.7% |  |  |
| Wavelet Frequency Refined Model |  |  |  |  |
| formula.crossed<-"Wavelet_Frequency ~ (1 \| Age) + (1 \| Sex) + (1 \| Mouse_Number) + (1 \| Position) + (1 \| ROI) + (1 \| Mouse_Number:Position) + (1 \| Mouse_Number:ROI) + (1 \| Position:ROI)" |  |  |  |  |
| Mouse number: ROI | 1.16 | 0.3% | n per group = 75 | n per group = 59 |
| Mouse number : position | 3.05 | 9.0% |  |  |
| Position: ROI | 0.00 | 0% |  |  |
| Mouse number | 2.83 e 1 | 82.9% |  |  |
| ROI | 0.00 | 0% |  |  |
| Position | 0.00 | 0% |  |  |
| Sex | 2.82 e -7 | <0.01% |  |  |
| Age | 2.10 | 6.2% |  |  |
| Residual | 5.53 e -1 | 1.6% |  |  |
| Wavelet Frequency SD Initial Model |  |  |  |  |
| lmer(Outcome ~ 1 + (1 \| Age) + (1 \| Sex) + (1 \| Mouse_Number) + (1 \| Position) + (1 \| ROI) + (1 \| Mouse_Number:Position) + (1 \| Mouse_Number:ROI) + (1 \| Position:ROI) + (1 \| Position:Sex) + (1 \| ROI:Sex) + (1 \| Position:Age) + (1 \| ROI:Age) + (1 \| Sex:Age)) |  |  |  |  |
| Mouse number: ROI | 5.64 e -1 | 7.8% | N/A | N/A |
| Mouse number : position | 1.64 | 22.6% |  |  |
| Position: ROI | 0.00 | 0.0% |  |  |
| ROI: Age | 0.00 | 0.0% |  |  |
| ROI: Sex | 0.00 | 0.0% |  |  |
| Mouse number | 6.73 e -1 | 9.3% |  |  |
| ROI | 2.26 e -1 | 3.1% |  |  |
| Position: Age | 0.00 | 0.0% |  |  |
| Position: Sex | 3.120 e -1 | 4.3% |  |  |
| Sex: Age | 0.00 | 0.0% |  |  |
| Position | 0.00 | 0.0% |  |  |
| Sex | 0.00 | 0.0% |  |  |
| Age | 1.70 e -9 | <0.01% |  |  |
| Residual | 3.85 | 53.0% |  |  |
| **Wavelet Frequency SD Refined Model** |  |  |  |  |
| formula.crossed <- "Wavelet_Frequency_SD ~ (1 \| Age) + (1 \| Sex) + (1 \| Mouse_Number) + (1 \| Position) + (1 \| ROI) + (1 \| Mouse_Number:Position) + (1 \| Mouse_Number:ROI) + (1 \| Position:ROI) + (1 \| Position:Sex)" |  |  |  |  |
| Mouse number: ROI | 7.06 | 4.5% | n per group = 1570 | n per group = 1298 |
| Mouse number : position | 5.48 | 34.6% |  |  |
| Position: ROI | 0.00 | 0.0% |  |  |
| Mouse number | 6.73 e -1 | 42.5% |  |  |
| ROI | 2.82 e -2 | 1.8% |  |  |
| Position | 0.00 | 0.0% |  |  |
| Sex | 4.48 e -10 | <0.01% |  |  |
| Position:Sex | 1.04 e -1 | 6.6% |  |  |
| Age | 6.28 e -10 | <0.01% |  |  |
| Residual | 1.61 e -1 | 10.1% |  |  |
| **Peak Valley Amplitude Initial Model** |  |  |  |  |
| lmer(Outcome ~ 1 + (1 \| Age) + (1 \| Sex) + (1 \| Mouse_Number) + (1 \| Position) + (1 \| ROI) + (1 \| Mouse_Number:Position) + (1 \| Mouse_Number:ROI) + (1 \| Position:ROI) + (1 \| Position:Sex) + (1 \| ROI:Sex) + (1 \| Position:Age) + (1 \| ROI:Age) + (1 \| Sex:Age)) |  |  |  |  |
| Mouse number: ROI | 3.47 e-5 | 5.9% | N/A | N/A |
| Mouse number : position | 1.56 e-4 | 26.4% |  |  |
| Position: ROI | 3.47 e-6 | 0.6% |  |  |
| ROI: Age | 2.91 e-6 | 0.5% |  |  |
| ROI: Sex | 2.40 e-6 | 0.4% |  |  |
| Mouse number | 1.02 e-4 | 17.4% |  |  |
| ROI | 0.00 | 0.0% |  |  |
| Position: Age | 0.00 | 0.0% |  |  |
| Position: Sex | 1.09 e-13 | <0.01% |  |  |
| Sex: Age | 5.96 e-13 | 25.1% |  |  |
| Position | 5.96 e-13 | <0.01% |  |  |
| Sex | 6.41 e-12 | <0.01% |  |  |
| Age | 4.46 e-5 | 7.6% |  |  |
| Residual | 9.57 e-5 | 16.2% |  |  |
| **Peak Valley Amplitude Refined Model** |  |  |  |  |
| formula.crossed<-"Peak_Valley_Amplitude ~ (1 \| Age) + (1 \| Sex) + (1 \| Mouse_Number) + (1 \| Position) + (1 \| ROI) + (1 \| Mouse_Number:Position) + (1 \| Mouse_Number:ROI) + (1 \| Position:ROI) + (1 \| Sex:Age)" |  |  |  |  |
| Mouse number: ROI | 4.718 e-6 | 1.3% | n per group = 19 | n per group = 15 |
| Mouse number : position | 5.189 e-5 | 14.6% |  |  |
| Position: ROI | 1.499 e-7 | <0.01% |  |  |
| Mouse number | 1.020 e -4 | 28.6% |  |  |
| ROI | 2.6124 e-7 | 0.1% |  |  |
| Sex:Age | 1.48 e -4 | 41.6% |  |  |
| Position | 0 | 0% |  |  |
| Sex | 3.148 e-8 | <0.01% |  |  |
| Age | 4.505 e-5 | 12.6% |  |  |
| Residual | 3.985 e-6 | 1.1% |  |  |
| Wavelet Amplitude Initial Model |  |  |  |  |
| lmer(Outcome ~ 1 + (1 \| Age) + (1 \| Sex) + (1 \| Mouse_Number) + (1 \| Position) + (1 \| ROI) + (1 \| Mouse_Number:Position) + (1 \| Mouse_Number:ROI) + (1 \| Position:ROI) + (1 \| Position:Sex) + (1 \| ROI:Sex) + (1 \| Position:Age) + (1 \| ROI:Age) + (1 \| Sex:Age)) |  |  |  |  |
| Mouse number: ROI | 2.73 e -5 | 6.1% | N/A | N/A |
| Mouse number : position | 1.04 e -4 | 23.4% |  |  |
| Position: ROI | 4.12 e -6 | 0.9% |  |  |
| ROI: Age | 0.00 | 0.0% |  |  |
| ROI: Sex | 2.05 e -6 | 0.5% |  |  |
| Mouse number | 7.00 e -5 | 15.8% |  |  |
| ROI | 1.10 e -14 | < 0.01% |  |  |
| Position: Age | 1.68 e -5 | 3.8% |  |  |
| Position: Sex | 2.51 e -13 | <0.01% |  |  |
| Sex: Age | 1.02 e -4 | 23.0% |  |  |
| Position | 3.39 e -14 | <0.01% |  |  |
| Sex | 0.00 | 0.0% |  |  |
| Age | 4.90 e -5 | 11.0% |  |  |
| Residual | 6.88 e -5 | 15.5% |  |  |
| Wavelet Amplitude Refined Model |  |  |  |  |
| formula.crossed <- "Wavelet_Amplitude ~ (1 \| Age) + (1 \| Sex) + (1 \| Mouse_Number) + (1 \| Position) + (1 \| ROI) + (1 \| Mouse_Number:Position) + (1 \| Mouse_Number:ROI) + (1 \| Position:ROI) + (1 \| Position:Age) + (1 \| Sex:Age)" |  |  |  |  |
| Mouse number: ROI | 3.61 e -6 | 1.3% | n per group = 17 | n per group = 13 |
| Mouse number : position | 3.47 e -5 | 13.0% |  |  |
| Position: ROI | 1.90 e -7 | 0.1% |  |  |
| Mouse number | 6.98 e -5 | 26.1% |  |  |
| ROI | 1.10 e -11 | <0.01% |  |  |
| Position: Age | 5.57 e -6 | 2.1% |  |  |
| Sex: Age | 1.02 e -4 | 38.2% |  |  |
| Position | 0.00 | 0% |  |  |
| Sex | 2.23 e -12 | <0.01% |  |  |
| Age | 4.86 e -5 | 18.2% |  |  |
| Residual | 2.86 e -6 | 1.1% |  |  |
| Mean Intensity Initial Model |  |  |  |  |
| lmer(Outcome ~ 1 + (1 \| Age) + (1 \| Sex) + (1 \| Mouse_Number) + (1 \| Position) + (1 \| ROI) + (1 \| Mouse_Number:Position) + (1 \| Mouse_Number:ROI) + (1 \| Position:ROI) + (1 \| Position:Sex) + |  |  |  |  |
| (1 \| ROI:Sex) + (1 \| Position:Age) + (1 \| ROI:Age) + (1 \| Sex:Age)) |  |  |  |  |
| Mouse number: ROI | 2.68 e -3 | 12.8% |  |  |
| Mouse number : position | 7.27 e -3 | 34.6% |  |  |
| Position: ROI | 6.22 e -4 | 3.0% |  |  |
| ROI: Age | 0.00 | 0.0% |  |  |
| ROI: Sex | 1.94 e 04 | 0.9% |  |  |
| Mouse number | 7.39 e -3 | 35.1% |  |  |
| ROI | 1.00 e -3 | 4.8% |  |  |
| Position: Age | 0.00 | 0.0% |  |  |
| Position: Sex | 2.51 e 12 | <0.01% |  |  |
| Sex: Age | 0.00 | 0.0% |  |  |
| Position | 1.94 e -12 | <0.01% |  |  |
| Sex | 3.05 e -12 | <0.01% |  |  |
| Age | 0.00 | 0.0% |  |  |
| Residual | 1.86 e -3 | 8.9% |  |  |
| Mean Intensity Refined Model formula.crossed<-"Mean_Intensity ~ (1 \|Age) + (1 \|Sex) + (1 \|Mouse_Number) + (1 \|Position) + (1 \|ROI) + (1 \|Mouse_Number:Position) + (1 \|Mouse_Number:ROI) + (1 \|Position:ROI)" |  |  |  |  |
| Mouse number: ROI | 3.51 e -4 | 3.4% | n per group = 485 | n per group = 393 |
| Mouse number : position | 2.42 e -3 | 23.3% |  |  |
| Position: ROI | 2.59 e -5 | 0.2% |  |  |
| Mouse number | 7.38 e -3 | 71.0% |  |  |
| ROI | 1.30 e -4 | 1.3% |  |  |
| Position | 0.00 | 0.0% |  |  |
| Sex | 0.00 | 0.0% |  |  |
| Age | 0.00 | 0.0% |  |  |
| Residual | 7.76 e -5 | 0.7% |  |  |

For power calculations, the n per group needed to detect a difference in outcome between age groups was calculated. Note that, strictly speaking, this is the number of truly independent measures required (when ICC = 0). Repeat measurements in a mouse are not independent measures due to clustering within animal. However, this calculation nonetheless gives a sense of how many total measurements per group are required.

**Supplemental Table 2:** Variance Components Analysis: Human Data.

| Random effects: | Variance | % attributable variance |
| --- | --- | --- |
| **Peak Valley Frequency** |  |  |
| Patient Number | 0.798 | 83.9% |
| ROI Number | 0.013 | 1.4% |
| Residual | 0.140 | 14.8% |
| **Wavelet Frequency** |  |  |
| Patient Number | 0.509 | 34.4% |
| ROI Number | 0.000 | 0.0% |
| Residual | 0.971 | 65.6% |
| **Wavelet Frequency SD** |  |  |
| Patient Number | 0.814 | 61.3% |
| ROI Number | 0.000 | 0.0% |
| Residual | 0.513 | 38.7% |
| **Peak Valley Amplitude** |  |  |
| Patient Number | 2.62 e -04 | 25.1% |
| ROI Number | 2.60 e -05 | 2.5% |
| Residual | 7.56 e -04 | 72.4% |
| **Wavelet Amplitude** |  |  |
| Patient Number | 7.46 e -05 | 33.3% |
| ROI Number | 1.64 e -05 | 7.3% |
| Residual | 1.33 e -04 | 59.4% |
| **Mean Intensity** |  |  |
| Patient Number | 0.00222 | 40.3% |
| ROI Number | 0.000 | 0.0% |
| Residual | 0.00329 | 59.7% |

**Model Used for Variance Components Analysis in Humans:** lmer(Outcome ~ 1 + (1|Patient\_Number) + (1|ROI\_Number))

